# Hypoxic conditions promote a proliferative, poorly differentiated, and pro-secretory phenotype in COPD lung tissue progenitor cells in vitro

**DOI:** 10.1101/2022.03.22.484879

**Authors:** Tina P. Dale, Michael D. Santer, Mohammed Haris, Wei Zuo, Nicholas R. Forsyth

## Abstract

Chronic obstructive pulmonary disease patients experience variable symptoms dependent on the presence of an emphysematous versus a chronic bronchitis phenotype. Both presentations can be associated with lung tissue and systemic hypoxia, at its most severe leading to Cor pulmonale. Despite this, minimal attention has been given to the effects of hypoxia at the cellular disease level.

We isolated and cultured progenitor cells from the distal lung tissue of a 64 year-old, male, emphysematous donor in ambient (21%) and hypoxic (2%) oxygen conditions. Proliferative capacity was determined on collagen coated culture plastic and growth-inactivated 3T3-J2 co-cultures. Epithelial (E-cadherin and pan-cytokeratin) and progenitor (TP63, cytokeratin 5) marker expression were examined. Expanded cells were differentiated at air-liquid interface and ciliated, mucous producing, and club cell populations identified.

Isolated cells were positive for the epithelial, pan-cytokeratin and E-cadherin, and progenitor, TP63 and cytokeratin 5, cell markers at isolation and again at passage 5. Passage 5 expanded cells in hypoxia had increased the proportion of TP63 expressing cells by 10% from 51.6 ± 1.2% to 62.6 ± 2.3% (p ≤ 0.01). Proliferative capacity was greater in 3T3J2 co-cultured cells overall and in 2% oxygen this supported the emergence of a proliferation unrestricted population with a limited differentiation capacity. Cells expanded on collagen I in either oxygen underwent differentiation having been expanded with the production of ciliated cells positive for βIV tubulin, and mucin 5ac, mucin 5b and CC10 positive secretory cells. Epithelial barrier formation was reduced significantly (p ≤ 0.0001) in hypoxia-expanded cells compared to normoxia. qRT-PCR showed higher expression of mucins in 2% expanded cells, significantly so with *MUC5B* (P ≤ 0.05) although mucin protein secretion was greater in 21% expanded cells.

Concomitantly these results demonstrate that hypoxia promotes a proliferative phenotype while reducing the overall differentiation capacity of the cells. Further, the retained differentiation potential becomes skewed to a more secretory phenotype demonstrating that hypoxia may be contributing to disease symptom and severity in COPD patients.

## Introduction

Chronic obstructive pulmonary disease (COPD) is the most common chronic respiratory disorder. Affecting around 12% of the global population^1^, it accounts for 55% of total respiratory disorder prevalence^2^ and as a result is the third leading cause of death worldwide, responsible for 6% of all deaths^3^. As a chronic disorder it is associated with considerable morbidity for affected individuals and presents a significant economic burden with an average annual per patient cost (direct and indirect) approaching $20000 in the UK and in excess of $30000 in the US^4^. Therapeutic options remain severely limited, with the management of symptoms and treatment of disease exacerbations as they arise continuing to be the norm. Despite its high prevalence and impact on global health, due to the complexity of the disease there is limited understanding of its etiology and pathology. Phenotypically the disease can present as either predominantly emphysematous, where alveolar tissue destruction is widespread; or as predominantly bronchitic, where a cough and excess sputum dominate; or in many cases with a substantial degree of overlap. Many patients also present with considerable co-morbidities, including other respiratory disorders such as asthma, further complicating study of the disease. Controlled *in vitro* research could provide invaluable insights into the causes, diagnosis, progression and treatment of COPD.

Excepting those individuals with a known alpha-1-antitrypsin deficiency (representing approximately 1% of COPD cases) the details of disease initiation, progression, and the reasons for such a diverse presentation, are not well defined^5^. COPD incidence is higher in males than females, increases with age and is strongly associated with smoking where prevalence is more than twice as high as in never-smokers^1^. Nevertheless, it remains unclear why some smokers develop COPD and others do not. Secondary to smoking, air pollution is a significant causative factor^6^. As a result to date there has been a strong research focus on the effects of cigarette smoke and the resultant inflammatory mediators in COPD.

Hypoxia is rare in healthy adult lung tissue but more common during fetal development and in diseased states, where hypoxia may be present in pockets of diseased tissue in the lungs dependent on disease severity^7^. The effects of COPD can have a profound effect on oxygenation in COPD patients, systemically this occurs as a result of a ventilation/perfusion (V/Q) mismatch in the lungs. All presentations of the disease can reduce the efficiency of oxygenation; emphysema reduces the available alveolar tissue for gas exchange generally causing a decrease in perfusion compared to ventilation and a high V/Q ratio. In contrast, airway disease is more likely to result in poor ventilation with an increase in perfusion to compensate, resulting in a low V/Q ratio. Exacerbations and even sleeping can serve to worsen the problem^8^. At its most severe, poor oxygenation can lead vasoconstriction within areas of the lung, causing pulmonary hypertension and potentially leading to right-sided heart failure (Cor pulmonale). More localized tissue hypoxia is also present with hypoxia inducible factor (HIF)-1α, a primary mediator of the hypoxic response, upregulated in association with bronchial tissue undergoing COPD-associated airway remodeling and associated goblet cell hyperplasia^9^. Despite this there is a scarcity of data regarding the direct impacts of hypoxia on lung cellular behaviour. The limited evidence available has shown a number of effects including increased oxidative stress, inflammatory markers, and pro-apoptotic genes in animal models^10–12^. Similarly cell-line models have also demonstrated increased oxidative stress, increased pro-inflammatory mediators, reduced anti-inflammatory mediators and reduced surfactant proteins^10^. Many effects of hypoxia can be induced both via continuous hypoxia or by hypoxia/reoxygenation events^12,11,10^, both of which can be found in COPD patients.

We have previously presented a protocol for the establishment of porcine airway epithelial cell cultures whereby with reduced oxygen conditions and Y-27632 RI supplementation cells underwent unlimited proliferation even in the absence of a feeder cell population^13^, a modification of a conditionally reprogrammed cell protocol that extended the proliferative span of epithelial cells, including airway cells^14,15^. However, concurrent with the enhanced proliferation we saw impaired differentiation. Herein we extend and transfer that protocol to human distal airway stem cells (hDASCs) obtained from a donor with COPD (emphysematous) expanded in normoxia (21% ambient oxygen) and hypoxia (2% oxygen). In contrast to our previous animal model, we noted that when cultured on a collagen coated substrate with RI, cells continued to undergo growth arrest, although proliferation was enhanced by a hypoxic environment. 3T3J2 feeder co-culture further enhanced proliferation, particularly in a hypoxic environment where proliferation appeared to be unlimited. In agreement with our porcine model prolonged expansion of cells impaired subsequent differentiation capacity which became unbalanced in favour of a pro-secretory phenotype.

## Materials and methods

### Cell culture

Human distal lung tissue was obtained from a 64 year old, male, emphysematous donor (FEV1 44% of predicted, FVC 48% of predicted, FEV1/FVC 29%) undergoing surgical lung resection at the Royal Stoke University Hospital with fully informed written consent and full Health Research Authority (HRA) ethical approval (16/WM/0447). Tissue was reviewed upon excision by an experienced Consultant Histopathologist to confirm the absence of unexpected gross abnormalities prior to same-day transfer to the cell culture laboratory on ice. Upon examination no cartilaginous airways were seen to be present in the tissue.

The tissue was washed in three changes of 500 μg/mL gentamicin sulphate (Thermo Scientific Alfa Aesar, UK), 100 μg/mL ciprofloxacin (Thermo Scientific, UK), 1000 IU/mL penicillin, 1 mg/mL streptomycin, and 2.5 μg/mL amphotericin B in Hank’s balanced salt solution (HBSS) (Lonza, UK). It was then finely minced and digested overnight on a rocker at 4 °C in 1 mg/mL protease XIV (Sigma-Aldrich, UK), 0.005% trypsin (Lonza, UK), and 10 ng/mL DNAse I (Sigma-Aldrich, UK) in F12 : DMEM (1 : 1) (Lonza, UK). Following digestion the suspension was passed through sterile gauze and a 70 µm cell strainer before centrifugation at 350 g for 10 minutes, red blood cells were lysed with RBC lysis buffer (Sigma-Aldrich, UK), and the cells counted with trypan blue (Sigma-Aldrich, UK) for dead cell discrimination. Cells were then seeded on tissue culture plastic pre-coated with a 10 μg/cm^2^ type 1 rat tail collagen solution (Corning, UK). Cells were incubated in cFAD+RI (DMEM : F12 (75 : 25), 10% (v/v) FBS (Biosera, UK), 1% (v/v) non-essential amino acids (NEAA), 2 mM L-glutamine (Lonza, UK), 24 μg/mL adenine, 5 μg/mL transferrin, 5 μg/mL insulin, 0.4 μg/mL hydrocortisone, 0.13 μg/mL triiodothyronine (all Sigma-Aldrich, UK), and 10 ng/mL epidermal growth factor (EGF) (Peprotech, UK), 10 µM Y-27632 dihydrochloride Rho-kinase inhibitor^13^ (Selleck Chemicals, UK) in either standard tissue culture incubators (21% O_2_ (air), 5% CO_2_, 5% humidity, 37 °C) or tri-gas incubators (2% O_2_, 5% CO_2_, 73% N_2_, 95% humidity, 37 °C) as indicated throughout. During routine culture media changes were performed twice weekly, when necessary cells were subcultured enzymatically using 0.1% (w/v) trypsin/0.04% (w/v) ethylenediaminetetraacetic acid (EDTA) (5 g/L trypsin and 2 g/L EDTA diluted 1 in 5 in calcium- and magnesium-free PBS) (Lonza, UK).

To produce feeder layers for co-culture 3T3J2 cells (Kerafast, US) were expanded (maximum of 12 additional passages from receipt) in 4.5 g/L DMEM supplemented with 10% iron-supplemented bovine calf serum (Seradigm, US), (1% (v/v) non-essential amino acids (NEAA), 2 mM L-glutamine (Lonza, UK), 100 IU/mL penicillin,100μg/mL streptomycin, and 0.25μg/mL amphotericin B. Media was changed twice per week and cells were passaged enzymatically as necessary with 0.05% (w/v) trypsin/0.02% (w/v) EDTA. Cells were inactivated by culture in the presence of 10 μg/mL of mitomycin C (Tocris, UK) in culture media for 2 hours, followed by 3 PBS washes, trypsinisation, and cryopreservation until required. Cells were recovered from cryopreservation at a density of 15000 cells/cm^2^ approximately 16 -24 hours before being needed.

### Cell proliferation

At early passage cell population doublings were estimated based on the split ratio used at passage. At later passage the population doublings were determined by cell counting and the use of the formula n = 3.32 (log UCY - log l, where n = the number of PD, UCY = the cell yield, l = the starting cell number.

### Flow cytometry

Flow cytometry was used to determine the forward scatter (FSC) and side scatter (SSC) of the cultured cells across passages/population doublings and TP63 expression at passage 5. For FSC and SSC analysis cells were trypsinised, washed with PBS (Lonza, UK), re-suspended in 400 µL of PBS and data obtained immediately on a Cytoflex flow cytometer (Beckman Coulter Life Sciences). For TP63 analysis cells were trypsinised and washed with PBS before fixation in 2% (w/v) chilled paraformaldehyde (PFA) in PBS for 10 minutes. PFA was removed and cells permeabilised in 0.1% Triton-X (Sigma-Aldrich, UK) in PBS for 15 minutes and blocked with 5% BSA (Thermo Scientific, UK) for 15 minutes. Directly conjugated (Alexa Fluor^®^ 647) primary TP63 antibody (Abcam, ab246728, UK) and isotype control antibody (Cell Signalling Technology, 3452S, UK) were diluted as per the primary antibody manufacturer’s recommendation for flow cytometry to 0.08 µg/mL in 0.05% Triton-X in PBS and cells were re-suspended in 100 µL of antibody or isotype control solution for 30 minutes before washing with PBS and re-suspending in flow cytometry buffer (0.5% (w/v) BSA, 2 mM EDTA (Lonza, UK) in PBS) for analysis, unstained cells were also included. Fluorescence data was collected on the APC-A channel and for all analyses a gating strategy was employed whereby firstly debris and dead cells were gated out of the data, followed by doublet exclusion using the FSC-height and FSC-area parameters. All other events were included in the analyses. Data were analysed using Cytexpert software (Beckman Coulter).

### Air-Liquid Interface Cultures

To induce differentiation, cells were cultured at the air-liquid interface on ThinCert cell culture inserts (0.34 μm area, 0.45 μm pore size) (Greiner, UK) in sterile 24-well plates. Cells were seeded at 2 × 10^5^ cells/cm^2^ in cFAD+RI medium and cultured for 3 days, then medium changed to 1:1 cFAD+RI:Pneumacult ALI (Stemcell Technologies, UK) for 2 days to ensure monolayers were 100% confluent. Culture medium was then removed and replaced with complete PneumaCult ALI medium in the basal compartment only. All differentiation was performed at 21% O_2_. Medium was changed twice per week for 21-28 days.

### TEER

Transepithelial electrical resistance was determined for differentiating ALI cultures using a Millicell ERS2 epithelial volt-ohm meter. To measure TEER, medium was removed from culture inserts (apical and basal at day 0 and basal only at days 7, 14 and 21) and replaced with HBSS with calcium and magnesium and the barrier resistance determined by inserting the probe across the apical and basal compartments. The HBSS was then aspirated from both compartments and Pneumacult ALI medium returned to the basal compartments of cultures only, to restore ALI. TEER was calculated by subtracting the resistance measurement of HBSS and insert only from ALI measurements and multiplying by the insert culture area to give an Ω·cm^2^ value.

### Immunocytochemistry

Cells were fixed in either 95% (w/v) chilled methanol (Thermo Fisher, UK) or 4% (w/v) paraformaldehyde (Sigma-Aldrich, UK) and stored under PBS at 4 °C until staining. Cells were permeabilised with 0.1% (w/v) Triton X for 10 minutes and then incubated with 5% (w/v) BSA for 1 hour at room temperature before overnight incubation with primary antibody diluted in PBS at 4° C (vimentin, 1 in 500 (ab92547); pan-cytokeratin, 1 in 100 (ab86734); alpha smooth muscle actin, 1 in 100 (ab5694); E-cadherin, 1 in 50 (ab15148); P63, 1 in 300 (ab124762); cytokeratin 5, 1 in 100 (ab52635); mucin 5AC, 1 in 200 (ab3649); beta IV tubulin, 1 in 500 (ab179509); mucin 5AC (Abcam, UK); mucin 5B, 1 in 50 (sc-21768) (Santa Cruz Biotechnology, US). Primary antibody was then removed, and cells were washed in 3 changes of PBS prior to incubation with secondary antibody (DyLight 488 anti-rabbit IgG (ab96881) or DyLight 594 anti-mouse IgG (ab96881) (Abcam, UK)) for 1 hour at room temperature in the dark. The secondary antibody was then removed, and cells were washed in 2 changes of PBS before incubation with DAPI (Sigma-Aldrich, UK) for 5 minutes. Cells were then washed once, covered with PBS and imaged on a Nikon Eclipse T1 fluorescence microscope with a Photometrix Prime Mono camera or an Olympus FV1200 confocal microscope.

### Alcian Blue-Periodic Acid Schiff (AB-PAS) Staining

PBS was removed from basal and apical chambers of fixed ALI cultures and the samples covered with 1% Alcian blue in 3% aqueous acetic acid for 30 minutes, washed with distilled water until water remained clear followed by 0.5% periodic acid solution for 5 minutes, washed with distilled water, and covered with Schiff reagent (all Sigma-Aldrich, UK) for 20 minutes. Schiff reagent was removed, and the samples were washed with lukewarm tap water for 5 minutes. Culture inserts were imaged submerged in distilled water.

### Mucin 5ac and Mucin 5B ELISA

To collect samples for mucin quantification ALI cultures were established as described for 28 days. Cell samples were taken at days 0, 21 and 28 by lysis with RIPA buffer. At days 7, 14, 21 and 28 the apical surfaces of cultures were washed with 50 μL of HBSS for 15 minutes to collect secreted mucins. Apical washes were combined for analysis over the full culture period. Lysates and apical washes were stored at -80 °C prior to analysis.

To prepare lysates for analysis they were first treated with DNase I at 1 μg/mL for 30 minutes on ice and centrifuged for 20 minutes at 1000 g at 4 ⍰C to remove insoluble impurities. Mucin 5AC and mucin 5B sandwich ELISAs (FineTest, China)) were carried out in accordance with manufacturer’s directions.

### Gene expression analysis

Samples were lysed and total RNA extracted using a Qiagen RNeasy Mini Kit. RLT lysis buffer supplemented with 10 µL/mL β-mercaptoethanol (Sigma-Aldrich, UK) was pipetted onto the samples and the cells disrupted by pipetting before transfer to a microcentrifuge tube and storage at -80 °C until extraction. RNA was extracted using the kit as per the manufacturer’s protocol. Eluted RNA was quantified with a NanoDrop 2000 spectrophotometer (Thermo Scientific, UK) and stored at −80°C until required.

Relative gene expression in undifferentiated cells and 21 day ALI differentiated monolayers was assessed using quantitative reverse transcription–polymerase chain reaction (qRT-PCR) with the SuperScript III Platinum SYBRGreen OneStep qPCR Kit (Life Technologies, UK) and an AriaMX real-time thermal cycler. 5 ng of RNA was used per reaction and samples’ relative expression levels of MUC5AC, MUC5B, CC10 and TP63 were normalised to GAPDH expression with expression in differentiated cells expressed relative to undifferentiated cells (-ΔΔCT).

## Results

### Cell isolation and expansion

Cell expansion characteristics were determined for collagen-cultured cells isolated from a human COPD lung tissue donor in both standard, air oxygen, 21% O_2_ culture conditions and in reduced, 2% O_2_ conditions. Cells with a predominantly epithelial morphology were obtained from culture in either condition (Fig. 1A). Upon continuous passaging cells in 2% O_2_ displayed a more uniform morphology while in 21% O_2_ cells would stratify into two distinct populations: higher density epithelial cell colonies with very visible cell junctions (outlined in Fig. 1A upper) surrounded by more loosely organized cells with less visible cell-cell contacts. Flow cytometry quantification of the FSC-A characteristic confirmed that 21% cultures consistently had a higher mean FSC-A than 2% cultures (14.5-25.6% higher, statistically significant at all passages measured) (Fig. 1B) indicating a larger cell size; FSC-A increased as expected as cells approached growth arrest.

**Figure 1.**
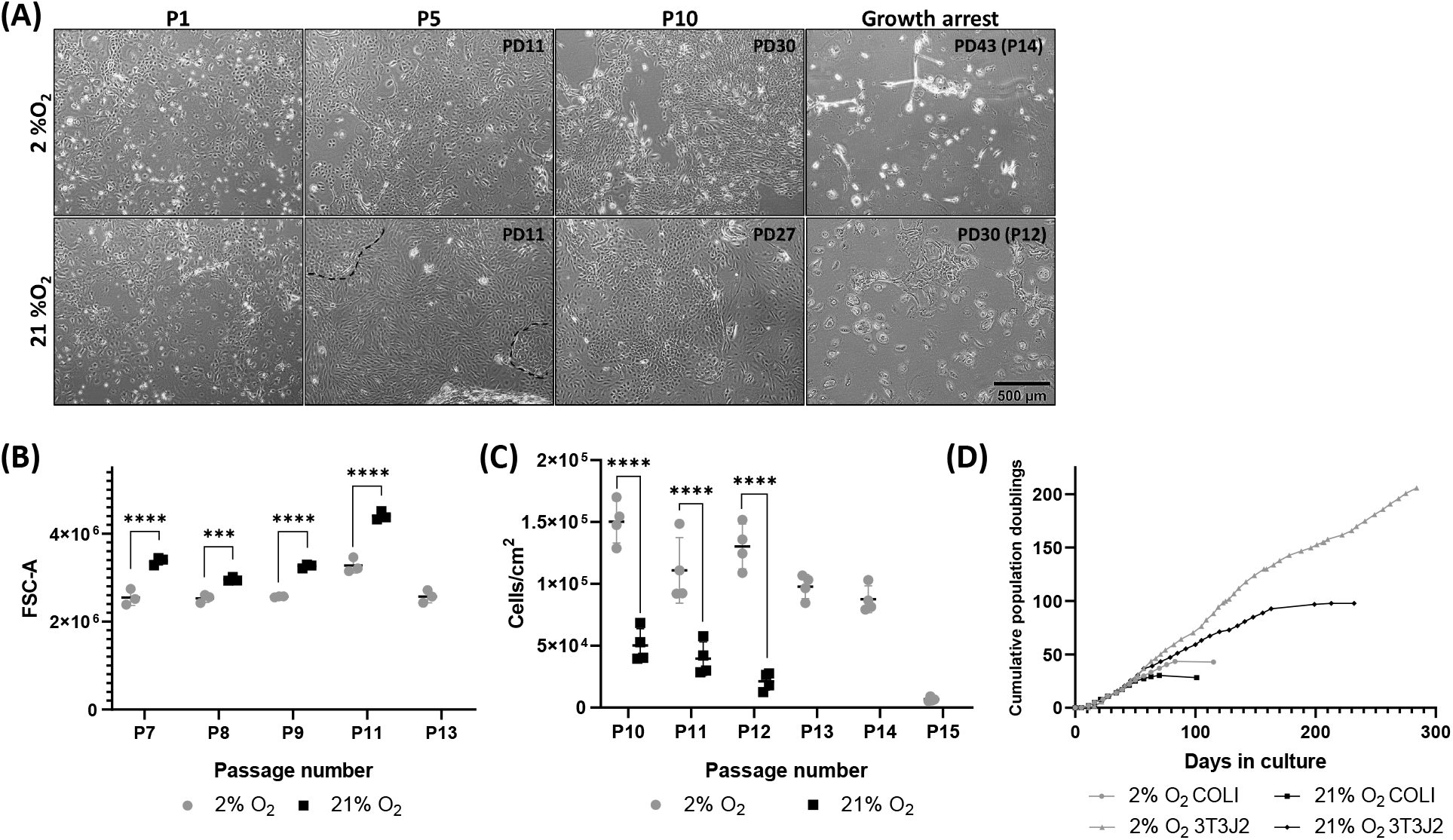
Reduced oxygen culture conditions increased total cell proliferation capacity and promoted a more uniform cell phenotype. (A) Phase contrast images of cells during expansion, acquired at passages 1, 5 and 10, and at the point of growth arrest in 2% (upper) or 21% (lower) O_2_ conditions, population doublings at passage are also indicated on the image. Cells in 21% O_2_ stratified into a looser epithelial phenotype with more compacted colonies scattered throughout (highlighted with a dashed line). Scale bar: 500 µm. (B) Mean flow cytometry FSC-A of cells to compare relative size at the passage numbers indicated. 21% cultured cells were consistently significantly larger than cells cultured at 2% O_2_. Due to growth arrest insufficient cells were available from 21% culture at P13 and beyond to compare 21% with 2% O_2_. Data are mean ± standard deviation, n=3. (C) Cell recovery from culture in 2% or 21% O_2_ at the passage numbers indicated, significantly higher cells/cm^2^ were recovered in 2% compared to 21% O_2_. Data are mean ± standard deviation, n=4. (D) Cumulative population doublings of cells cultured in 2% or 21% O_2_ on surfaces coated with a type I collagen substrate or with mitomycin C inactivated 3T3J2 feeder cells. On either substrate 2% O_2_ culture increased the proliferative capacity of the cells. Culture with 3T3J2 feeder cells also greatly increased the proliferative capacity of cells compared to collagen substrates, with 2% 3T3J2 cultures apparently immortalized. ***p≤0.001 ****p≤0.0001.

Cell counting at later passages confirmed that significantly higher cell numbers could consistently be recovered from 2% cultures than 21% cultures (Fig. 1C). Total proliferative capacity of the cells was assessed by estimating cumulative population doublings (Fig. 1D), with cells cultured in 21% achieving 30 PDs in 14 passages, in comparison 2% cultures were 43% higher, with 43 PDs from 14 passages. Unlike our previous porcine cultures^13^, neither condition supported unlimited proliferation.

Cells recovered and expanded on inactivated 3T3J2 feeder cells demonstrated increased proliferation in both oxygen conditions, with 21% cultured cells reaching 97 PDs in 200 days before growth arrest. In contrast, 2% cultured cells reached 206 PDs in 284 days with no sign of growth arrest (Fig. 1E). However, initial attempts to differentiate these significantly expanded cells resulted in poor monolayers (Fig. 2). These monolayers had only patchy cell coverage as indicated by broken regions of DAPI staining (Fig. 2A, 2B, 2C left panel) and a complete absence of ciliation, with β IV tubulin staining being localized cytoplasmically (Fig. 2A). Notably, despite this there was still significant staining for mucin 5AC (Fig. 2A), mucin 5B (Fig. 2B) and CC10 (Fig. 2C), the presence of mucin secretion was further confirmed by positive AB-PAS staining where cells were present (Fig. 1D). As a result of this poor differentiation the remaining data herein is focused on early passage cells expanded on collagen coated tissue culture surface.

**Figure 2.**
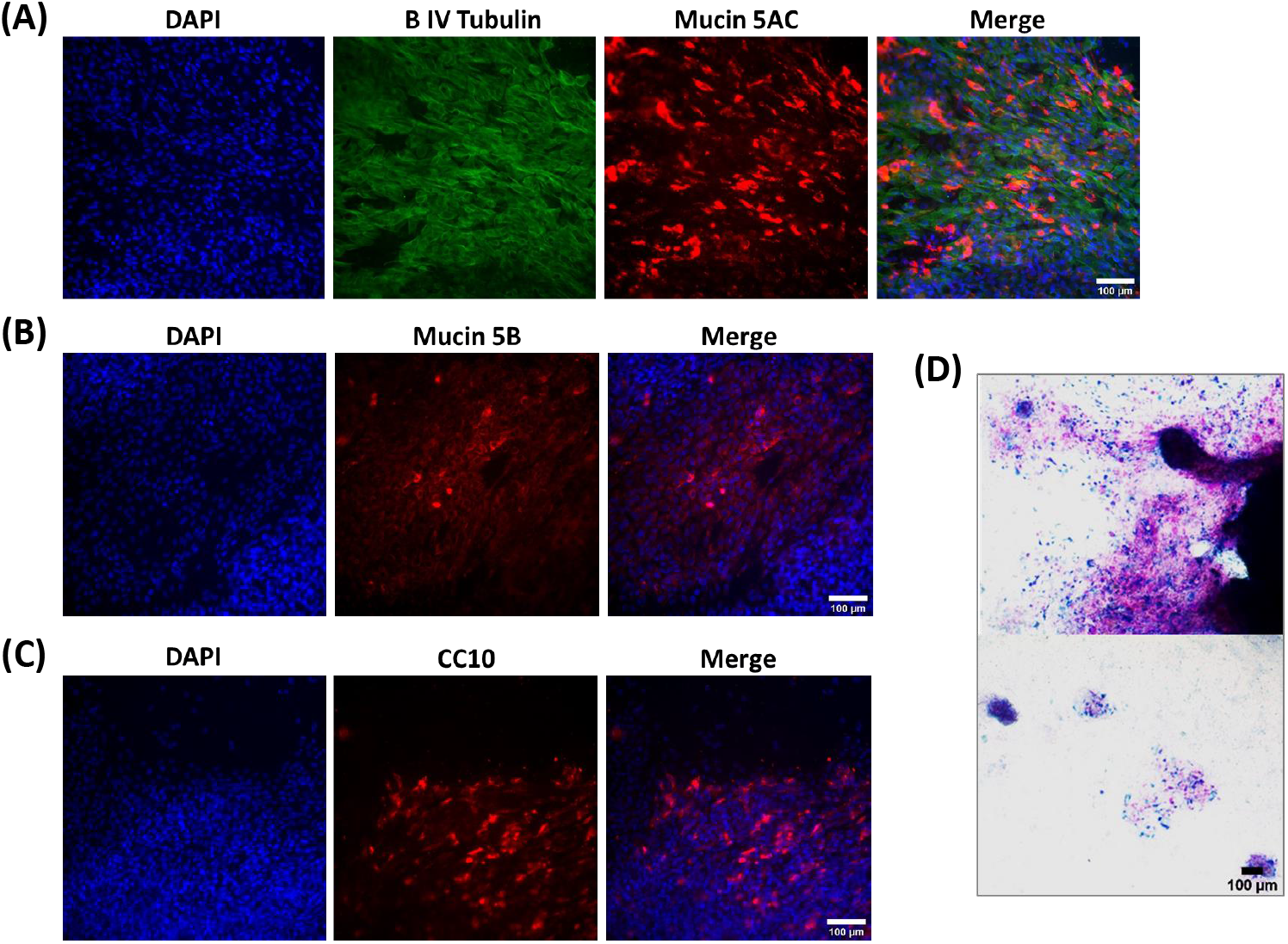
Lung progenitor cells highly expanded in hypoxia in the presence of 3T3J2 feeder cells differentiate poorly, lack cilia, and favour a pro-secretory phenotype when differentiated at ALI. (A) Immunofluorescence images of cells demonstrating cytoplasmic localization of β IV tubulin, and positive immunofluorescence staining for mucin 5AC and (B) mucin 5B. (C) Cells also exhibited positive expression of the club cell marker CC10. (D) Alcian blue and periodic acid Schiff histological staining confirmed mucin was produced in regions where the monolayer remained intact. Scale bar: 100 µm. Immunofluorescence images are counterstained with DAPI to identify nuclei.

### Cell characterization during recovery and expansion

Isolated cells at passage 0 were stained for the epithelial cell markers pan cytokeratin and E-cadherin (Fig 3A, 3B), and vimentin and smooth muscle actin (Fig. 2A, 2C) to establish whether initial cultures were predominantly epithelial or were contaminated with fibroblasts or smooth muscle cells. Cells stained positively with Pan cytokeratin and E-cadherin confirming their epithelial origin, positive staining with vimentin and smooth muscle actin was rare in either oxygen condition confirming high epithelial cell selection with our growth conditions.

**Figure 3.**
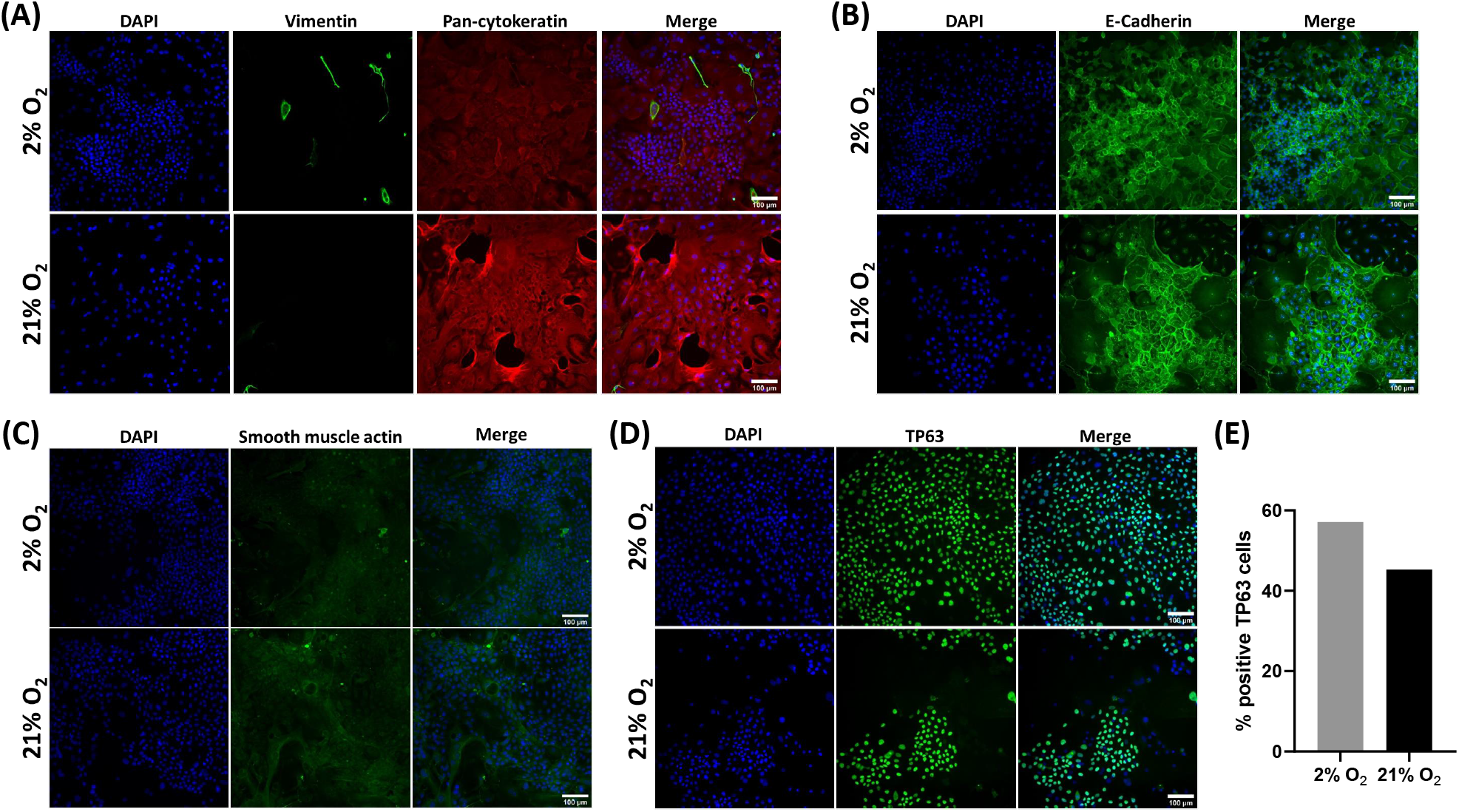
Recovered cells were characteristic of epithelial progenitor cells. Immunofluorescence images of cells at recovery (P0). Cells had widespread expression of the epithelial markers pan-cytokeratin (A) and E-cadherin (B) and minimal expression of vimentin (A) or smooth muscle actin (C). Cells were also frequently, but not universally, positive for the basal cell marker TP63 (D). Scale bar: 100 µm. Immunofluorescence images are counterstained with DAPI to identify nuclei. (E) Quantification of TP63 expression via image analysis showed that cells were more frequently positive for TP63 in 2% O_2_ than in 21% O_2_.

Cultures were then further stained with a TP63 antibody to confirm a progenitor/stem cell identity^16^. Positive reactivity was seen throughout the cultures with nuclear localization, excepting where cells were dividing. As too few cells were available at recovery to perform quantitative analysis via flow cytometry, TP63 expression in recovered cells was semi-quantified via image analysis using the Fiji distribution of ImageJ^17^ to determine the percentage of positive TP63 cells (Fig. 3E). As cultures were unevenly distributed across the wells, the whole-well field was imaged as 4 quadrant regions to avoid bias. In the 2% culture 57% of 2.04×10^4^ cells were TP63 positive compared to 45% of 5.48×10^3^ cells in 21% O_2_ culture.

Expression of epithelial, fibroblastic, and stem cell markers were further assessed in expanded cells at passage 5 (Fig. 4). Widespread expression of pan cytokeratin (Fig. 4A) and E-cadherin (Fig. 4B) was retained. Vimentin expression became more widespread (Fig. 4C) than at recovery but was concomitant with continued epithelial marker expression and the maintenance of an epithelial morphology, confirming that epithelial cells were expressing vimentin, rather than the presence of contaminating fibroblasts within the culture. TP63 expression (Fig. 4D) remained widespread, and as with passage 0, tended to be found within colonies of TP63 expressing cells. Along with TP63 expression we also confirmed positive expression of the progenitor cell-associated marker cytokeratin 5^18^ (Fig. 4E).

**Figure 4.**
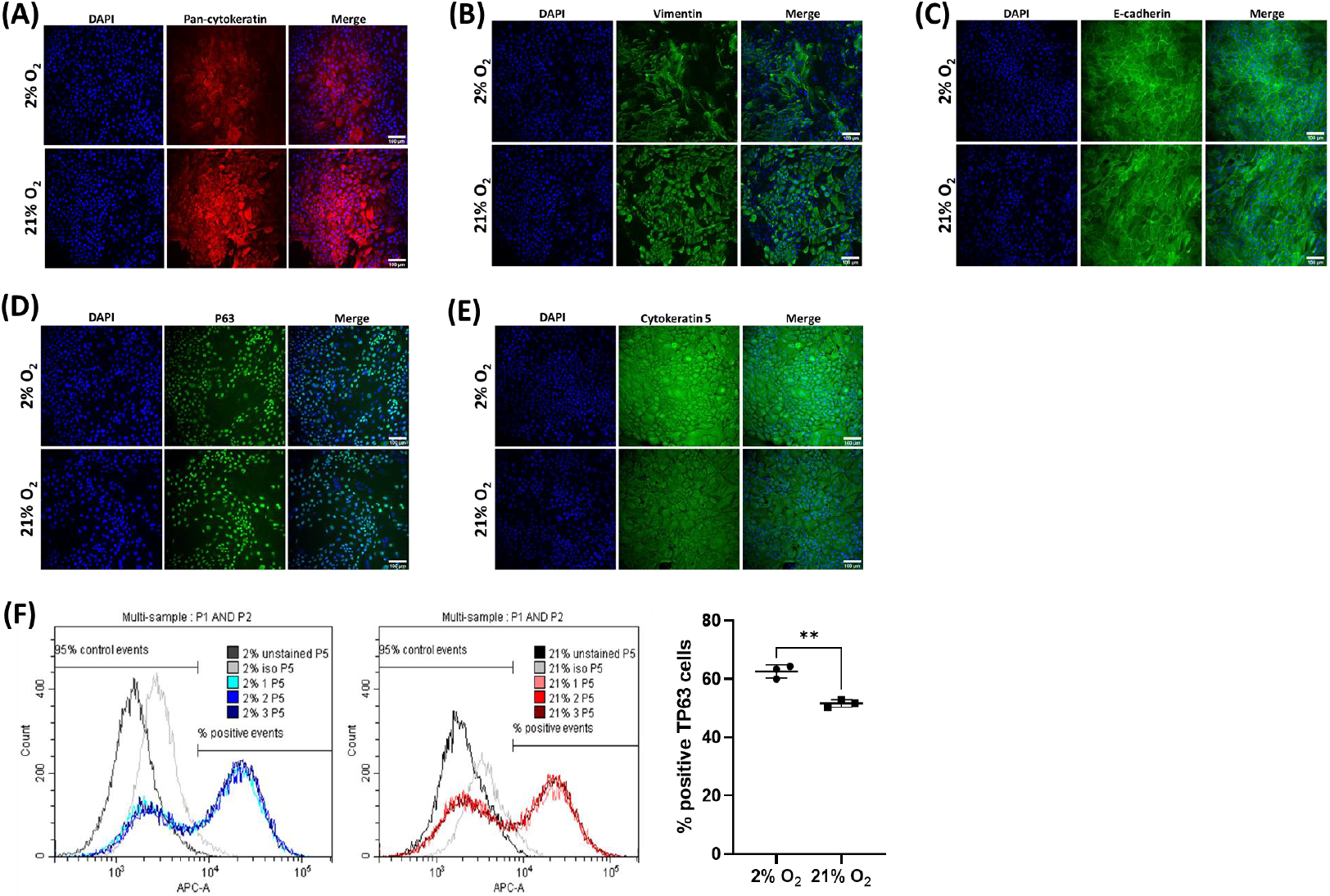
Culture expanded cells continue to express characteristic epithelial, basal cell markers. Immunofluorescence images of cells expanded to passage 5. Cells had continued expression of the epithelial marker Pan-cytokeratin (A), increased expression of vimentin (B) and positive expression of E-cadherin (C). Cells also remained positive for the basal cell marker TP63 (D) and had widespread expression of cytokeratin 5 (E). Scale bar: 100 µm. Immunofluorescence images are counterstained with DAPI to identify nuclei. (D) TP63 expression was quantified using flow cytometry with cells from 2% O_2_ cultures having significantly higher percent positive TP63 expression than cells cultured in 21% O_2_. Data are mean ± standard deviation, n=3, **p≤0.01.

TP63 expression at P0 appeared to be differential between the 2% and 21%, and expression was further quantified by flow cytometry. Flow quantification demonstrated a clear distinction between the negative and positive TP63 expressing populations in both oxygen conditions (Fig. 4F left and middle panels). There was significantly different TP63 expression between the oxygen levels at 62.6 ± 2.3% positive in 2% O_2_ compared to 51.6 ± 1.2% positive in 21% O_2_ (P≤0.01) (Fig. 4F right panel), similar levels to those determined from image analysis at P0.

### Cell differentiation

Cells were differentiated at ALI with a commercial differentiation medium for 21 days in a 21% O_2_ environment. Motile cilia developed on the surface of cultures and were visible using light microscopy. Immunocytochemistry confirmed mucociliary differentiation with expression of βIV tubulin (cilia) apically, mucin 5AC (Fig. 5A) and mucin 5B (Fig. 5B). Widespread staining was also present for the club cell marker CC10 (Fig. 5C). Histological staining with AB-PAS confirmed extensive mucous production across the culture surface (Fig. 5D).

**Figure 5.**
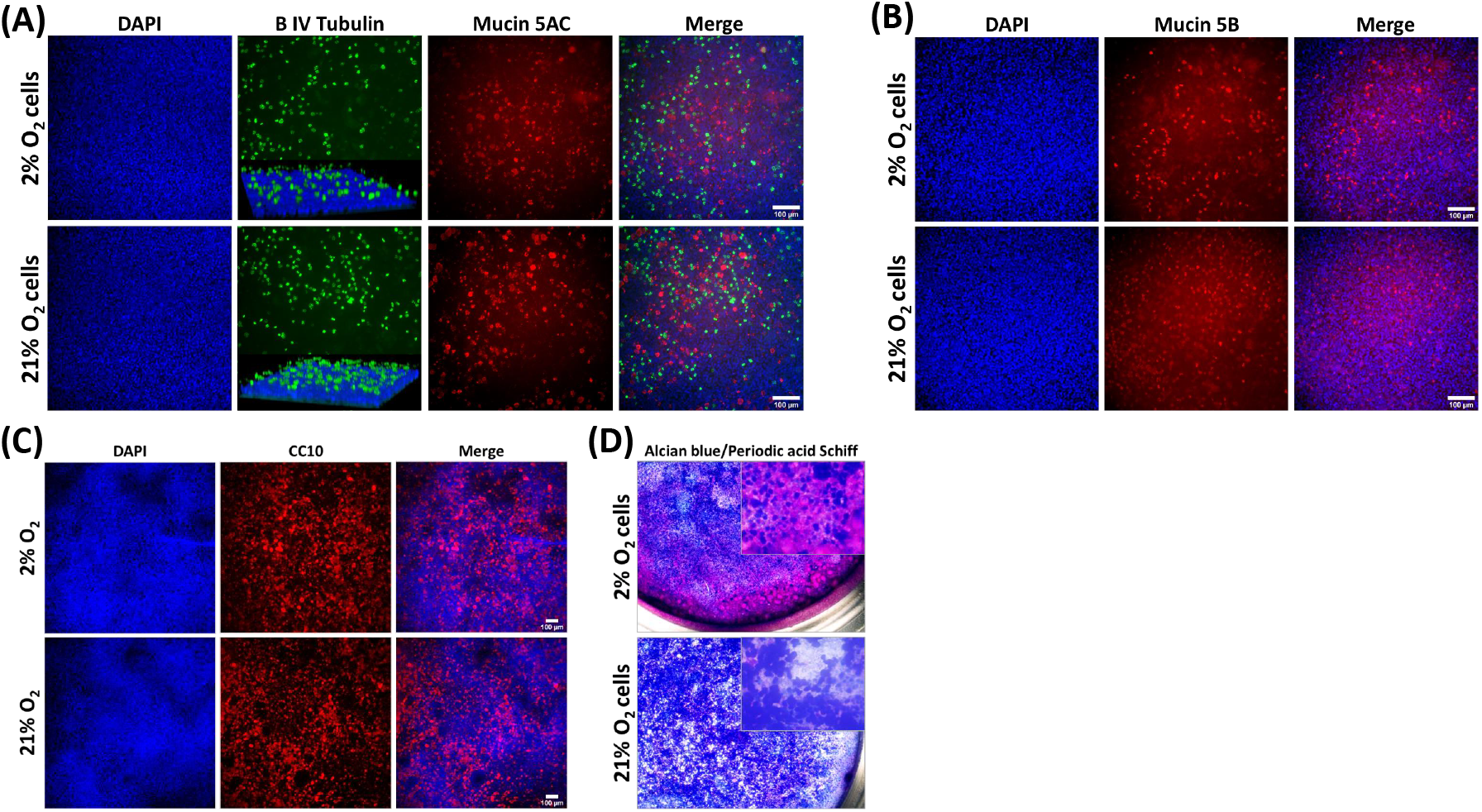
Cells expanded in either O_2_ condition underwent mucociliary differentiation when cultured in 21% O_2_ at an air-liquid interface. (A) Immunofluorescence images of cells demonstrating positivity for β IV tubulin (green) localized apically (inset is a three dimensional confocal microscope image), Mucin 5AC (red) and (B) Mucin 5B (red). (C) Cells also had widespread expression of the club cell marker CC10 in both O_2_ conditions. (D) Alcian blue and Periodic acid Schiff histological staining confirmed large amounts of mucin was produced in both culture conditions. Scale bar: 100 µm. Immunofluorescence images are counterstained with DAPI to identify nuclei.

The levels of differentiation markers *MUC5AC, MUC5B* and *CC10* were quantitatively examined at the gene expression level, along with *TP63* expression. Expression of the cilia marker β IV tubulin was not included as we have commonly observed its widespread expression in the cytoplasm of cells in poorly differentiated cultures (Fig. 2A), therefore an increase in message cannot be directly correlated with successful differentiation. Both mucins were significantly upregulated in comparison to their undifferentiated counterparts (Fig. 6A, 6B) with higher expression in 2% expanded cells compared to 21% which reached statistical significance (P≤0.05) for *MUC5B*. Similarly, the club cell marker CC10 was also significantly upregulated by differentiation (Fig. 6C). Interestingly, expression of the progenitor cell marker *TP63* (Fig. 6D) was not seen to decrease as a result of the differentiation process. 21% O_2_ expanded cells had a small and insignificant increase over expression levels in undifferentiated cells following differentiation at 21%, but 2% expanded cell cultures had a moderate and statistically significant (p≤0.001) upregulation in expression compared to their undifferentiated counterparts. This therefore resulted in a significant difference between the 2% and 21% expanded cell groups following differentiation (P≤0.05).

**Figure 6.**
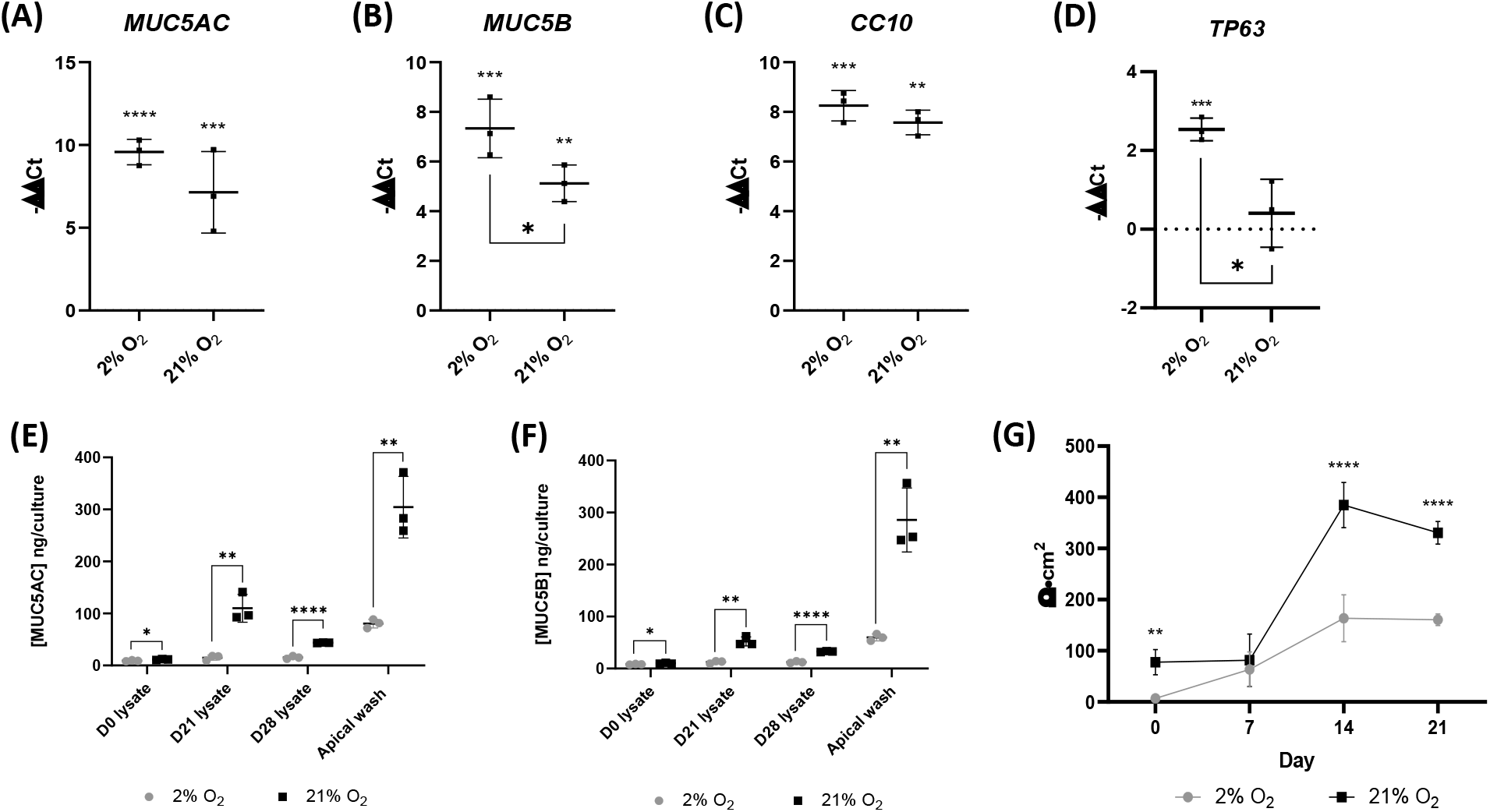
Cells expanded in 2% O_2_ conditions upregulate TP63 expression and demonstrate reduced differentiation with reduced mucous production and lower trans-epithelial electrical resistance (TEER). (A-D) qRT-PCR gene expression analysis for *MUC5AC, MUC5B, CC10* and *TP63*. Graphs show upregulation (-ΔΔCt) normalized to *GAPDH* expression and expressed relative to matched, undifferentiated cells. Significance indicators above are relative to undifferentiated cells, significance indicated below shows differences at day 21 between oxygen conditions. (A) *MUC5AC* expression was significantly upregulated in cells following 21 day differentiation at ALI. (B) *MUC5B* expression was significantly upregulated in cells following 21 day differentiation at ALI, expression was also significantly higher in 2% expanded cells than 21% expanded cells following differentiation. (C) *CC10* expression was significantly upregulated in cells following 21 day differentiation at ALI. (D) *TP63* expression was significantly upregulated in cells following 21 day differentiation at ALI, expression was significantly higher in 2% expanded cells than 21% expanded cells following differentiation. (E) Mucin 5AC protein produced during ALI differentiation over 28 days determined by ELISA. Significantly more Mucin 5AC was present in apical washes and lysates of differentiated cultures of 21% O_2_ expanded cells at all time points. (F) Mucin 5B protein produced during ALI differentiation over 28 days determined by ELISA. Significantly more Mucin 5B was present in apical washes and lysates of differentiated cultures of 21% O_2_ expanded cells at all time points. (G) TEER of differentiating cells at days 0, 7, 14 and 21. TEER was significantly higher in differentiating cultures of 21% O2 expanded cells at days 0, 14 and 21. Data are mean ± standard deviation, n=3 for qRT-PCR and Mucin ELISA n=6 for TEER, *p≤0.05 **p≤0.01 ***p≤0.001 ****≤0.0001

As we had observed differential expression of mucins at the gene level we quantified secreted and cell-associated mucin protein. In contrast to the gene-level data we found that levels of both mucin 5AC (Fig. 6E) and mucin 5B (Fig. 6F) whether secreted and measured by apical wash, or determined in the cell lysate, were significantly higher at all time points in 21% expanded cells than in 2% expanded cells following differentiation at 21% O_2_.

Finally, we determined the barrier function of the monolayers by measuring the TEER of every 7 days during the differentiation process (Fig. 6G). Monolayers expanded in 21% O_2_ developed a significantly higher TEER during differentiation than 2% O_2_ expanded cell monolayers (P≤0.0001), at more than double the Ω·cm^2^ value at day 14 (2% 163.5 ± 45.7 Ω·cm^2^, 21% 384.7 ± 44.3 Ω·cm^2^) and at day 21 (2% 160.6 ± 11.1 Ω·cm^2^, 21% 330.4 ± 22.5 Ω·cm^2^).

## Discussion

Over the past two decades there has been a stepwise improvement in knowledge that has supported the development of biologically relevant *in vitro* lung models to the point where these can now be considered representative of both healthy and diseased tissue. Nevertheless challenges remain, particularly surrounding the acquisition of sufficient cell numbers for both research and potential cell-based clinical therapies, and so far the investigation of disease etiology and progression remains in its infancy. Here we have described a technique to isolate and culture-expand TP63/cytokeratin 5 positive progenitor cells from a COPD patient. We have compared the proliferation and *in vitro* differentiation capacity of the cells following expansion in standard culture conditions (21% O_2_ ambient air incubator) and reduced oxygen culture conditions (2% O_2_ tri-gas controlled incubator), as the impact of hypoxia on lung cells has been subjected to limited investigation despite its prevalence both in lung tissues specifically, and systemically, in individuals with respiratory disorders. Overall, we noted an increased proliferative capacity in cells expanded in reduced oxygen, with culture in 2% O_2_, supplementation with Y-27632 and growth on 3T3J2 mitotically inactivated feeders resulting in an immortalized cell-line. However, we concomitantly saw an altered differentiation capacity where 2% O_2_ expansion of cells prior to differentiation in ALI conditions at 21% O_2_ resulted in reduced differentiation, with markedly reduced mucin production and barrier function. Overall, our results suggest that hypoxia exposure promotes a proliferative phenotype and concurrently impaired differentiation in these cells, even when increased oxygen availability is restored.

Reduced oxygen can lead to both pro- and anti-proliferative responses dictated by cell-type and circumstance^19^. In particular stem cell proliferation and maintenance of the undifferentiated state is maintained via multiple signals from the stem cell niche, including those generated by reduced oxygen conditions^20–23^. Similarly the impact on differentiation of hypoxia related signaling is likely to be widespread and variable. Mouse embryonic stem cell differentiation towards respiratory lineage cells can be promoted by hypoxia but is sensitive to both the extent and the duration of hypoxia^24^ and both human and mouse lung and airway progenitor cell fate is influenced by hypoxia with both potentially protective and pathological changes^25,26^.

The *in vitro* growth of cells from the lungs is reliant on the expansion of a progenitor cell population; a relatively large number of candidate cell-types have been identified, and similarly there are numerous prospective markers for these cells. It is often unclear whether different studies are referring to overlapping cell identities, and their differentiation/trans-differentiation capabilities, particularly where there is pathology present, have yet to be fully elucidated. *In vivo* these populations are responsible for tissue repair, maintaining cellular homeostasis and thus function. Typically, the progenitor cells are TP63 and cytokeratin 5 positive, this combination identifies progenitor cell types from both the proximal (generally referred to as basal cells) and more distal airways with further variation occurring in sub-populations of these cells^27,28^. We observed that while we had widespread cytokeratin 5 positivity, only a sub-set of cells was also positive for TP63 demonstrating culture heterogeneity. The TP63 positive cell fraction was larger, both at isolation, and following expansion in 2% O_2_. Zhao *et al* have described a cytokeratin 5 positive/TP63 negative population arising from double positive cells following engraftment in a bleomycin-injured mouse lung model^29^ suggesting that reduced oxygen may be inhibiting lineage progression in our cultures. Inhibition of the differentiation of cytokeratin 5 progenitors consequently impairs alveolar epithelial barrier formation^29^, something that we also observed with reduced ALI TEER in 2% cultured cells. The TP63 antibody we used reacts with the TAp63α full length, transactivating, isoform variants of TP63, there are reports that this isoform is not the dominant one in the human lung^30^; however, both isoforms have been identified in lung tissue samples at both the gene and protein level^31,32^. In mouse studies of DASCs the Tap63α isoform has been undetectable until 7 days post-infection with influenza virus, rapidly becoming more numerous in areas of damaged lung and decreasing following disease resolution suggesting that it is associated with a damage resolution response^33^. Interestingly we also observed an upregulation of TP63 in reduced oxygen suggesting a similar response in our human lung progenitors. However; subsequent differentiation of these cells at restored ambient oxygen remained poor in comparison to cells cultured in standard oxygen conditions, with sustained upregulation of TP63 suggestive of an unresolved response to the hypoxic insult.

While there is increasing evidence that a TP63+/KRT5+ cell population contributes to the healing response of the damaged lung ^18,34–38^ the presence of aberrant progenitor populations and altered differentiation in these damage-responsive cells may also be contributing significantly to changes associated with COPD. Rao *et al* describe a clone-dependent variation in the differentiation behavior of cells from COPD and normal lung. COPD lung yields greater total numbers of TP63+/KRT5+ proliferative clones with a much greater tendency towards aberrant differentiation, including goblet cell metaplasia, squamous cell metaplasia and pro-inflammatory squamous cell metaplasia; all typical of observations of COPD lungs^39^. Upregulation of both isoforms of TP63 has been associated with basal cell hyperplasia and abnormal bronchiolization in idiopathic pulmonary fibrosis^28,31,32^. Interestingly, TP63 has also been implicated in an epithelial to mesenchymal transition (EMT) response to hypoxia in cancer cells, including lung cancer^40^.

A significant contributor to the pathology of COPD, particularly where chronic bronchitis is a factor, is the presence of altered mucus production encompassing excess mucous secretion as a result of multiple factors, including goblet cell metaplasia/hyperplasia and hypertrophy, and hypersecretion^41^ changes in the mucin content of the mucous and altered mucous hydration. These factors modify the biophysical properties of mucous and contribute, alongside altered ciliary beat frequency, to impaired mucociliary clearance efficiency^42^. Together this can result in the development of airway mucous plugs and lead to airway collapse^43^. COPD patients suffering with chronic bronchitis are more likely to experience frequent exacerbations, hospitalization, and higher mortality^44^. The role of hypoxia in these changes has received little interrogation despite its prevalence in lung disorders, including COPD. Typically in the healthy lung mucin 5B is the predominant mucin at approximately 10 times higher levels than mucin 5AC in healthy individuals^45^. Mucin 5AC, while remaining the lesser mucin, proportionally increases in quantity with progression from the distal to more proximal airways^46^. In COPD patients the concentration of both mucins can be elevated, with the increase in mucin 5AC being disproportionately higher^47^. Interestingly, despite our ALI differentiation being induced in 21% oxygen, at the gene level we nevertheless observed a trend for increased expression of mucin 5AC and a significant increase in mucin 5B in cells previously expanded in reduced oxygen. It has previously been observed in cell line-based studies^48,49^ that in keeping with the presence of a hypoxia response element (HRE) hypoxia can directly induce mucin 5AC via hypoxia inducible factor (HIF) signaling^48^, our results where we see a sustained impact of hypoxia on later normoxic differentiation suggest that an additional mechanism, perhaps involving epigenetic changes, may be contributing to longer-term alterations in differentiation capacity. Similar responses could be active in COPD patients whereby intermittent periods of hypoxia, such as those experienced during exercise or infection/exacerbation lead to longer term deleterious changes.

Club cells, as identified herein by robust CC10 expression, were common in both 2% and 21% differentiated cultures and are largely absent in proximal airways^50^ confirming that we have retained a distal, bronchiolar airway phenotype during expansion and differentiation. Interestingly co-expression of mucins and CC10 has been observed in airways, again more common distally, although these cells were described as CC10 goblet cells^50^

In conclusion, while hypoxia increased the proliferative capacity of our cells it also impaired their subsequent differentiation capacity. After extensive expansion in hypoxia, ciliation could be lost completely while differentiation down secretory pathways with CC10, mucin 5AC and mucin 5B positive cells was disproportionally retained leading to an imbalance favouring excessive secretions. The effect of this imbalance has significant implications for epithelial repair and mucociliary clearance efficiency and therefore the biological alterations and symptoms seen in COPD patients.

## Acknowledgements

The authors would like to acknowledge the support of the UHNM theatre staff, Dr Sana Iftikhar and Dr Daniel Gey van Pittius for their assistance in acquiring participant lung tissue. This work was supported by funding from the North Staffordshire Medical Institute 50th Anniversary Award and The Royal Society International Exchange grant.

